# Application of single nuclei RNA sequencing to assess the hepatic effects of 2,3,7,8-tetrachlorodibenzo-*p*-dioxin

**DOI:** 10.1101/2020.04.07.030478

**Authors:** Rance Nault, Kelly A. Fader, Sudin Bhattacharya, Tim R. Zacharewski

**Affiliations:** Institute for Integrative Toxicology, Michigan State University, East Lansing, MI; Biochemistry & Molecular Biology, Michigan State University, East Lansing, MI; Biomedical Engineering, Pharmacology & Toxicology, Institute for Quantitative Health Science and Engineering, Michigan State University, East Lansing, MI, USA

**Keywords:** Single nuclei RNAseq, Liver, TCDD, Hepatotoxicity, NAFLD

## Abstract

Cell-specific transcriptional responses are lost in the averages of bulk RNA sequencing. We performed single nuclei RNA sequencing (snSeq) on frozen liver samples from male C57BL/6 mice in response to 2,3,7,8-tetrachlorodibenzo-*p*-dioxin (TCDD). Approximately 19,907 hepatic genes were detected across 16,015 sequenced nuclei from control and treated samples. Eleven cell-(sub)types were identified including distinct hepatocyte sub-populations, consistent with the cell diversity of the liver. TCDD increased macrophages from 0.5% to 24.7%, while neutrophils were only present in treated samples. The number of differentially expressed genes correlated with the basal expression level of *Ahr*. In addition to expected functional enrichments within each cell-(sub)type, RAS signaling was enriched in nonparenchymal cells. snSeq also identified a Kupffer cell subtype highly expressing *Gpnmb*, consistent with a dietary NASH model. Overall, snSeq distinguished cell-specific transcriptional changes and population shifts consistent with the hepatotoxicity of TCDD.

## INTRODUCTION

The liver is particularly susceptible to toxicity due to its close association with the gastrointestinal tract and xenobiotic metabolism capacity (Gu and Manautou, 2012). Liver toxicity is the primary driver of drug candidate attrition and a common cause of market withdrawal (Hornberg *et al.*, 2014). Similarly, environmental contaminant exposure is implicated in liver damage in humans, evident by increased levels of the liver damage biomarker alanine aminotransferase (ALT) in epidemilogical studies (Cave *et al.*, 2010; Heindel *et al.*, 2017; Kaiser *et al.*, 2012; Yorita Christensen *et al.*, 2013). Moreover, environmental contaminants such as 2,3,7,8-tetrachlorodibenzo-*p*-dioxin (TCDD), are potential contributing factors in the etiology of complex metabolic diseases such as obesity, type II diabetes, and non-alcoholic fatty liver disease (Cave *et al.*, 2010; *Heindel et al., 2017*; Kaiser *et al.*, 2012; Yorita Christensen *et al.*, 2013). For example, aryl hydrocarbon receptor (AhR) agonists, such as TCDD and related compounds, promote hepatic lipid accumulation (steatosis) and its progression to steatohepatitis (NASH) with fibrosis in mice (Fader *et al.*, 2017a; Nault *et al.*, 2016; Pierre *et al.*, 2014), while epidemiological studies suggest an association with metabolic disease in humans (Henriksen *et al.*, 1997; Taylor *et al.*, 2013).

Although bulk “omic” strategies have uncovered important knowledge on the mechanisms of hepatotoxicants, key molecular events, as well as the significance of temporal, spatial, and cellular heterogeneity, remain poorly understood for several chemicals. Single cell transcriptomic technologies provide the opportunity to investigate the transcriptomic responses to exogenous agents while also considering cellular heterogeneity and putative cell-cell interactions (Browaeys *et al.*, 2020). Single cell RNA sequencing (scSeq) can query the transcriptome at unprecendented resolution, characterizing rare cell types and developmental processes (Habib *et al.*, 2016; Su *et al.*, 2017), and further elucidating the significance of tissue spatial organization (Halpern *et al.*, 2018; Halpern *et al.*, 2017). scSeq analysis of a diet-induced NASH model revealed the expansion of a novel Kupffer cell subtype termed NASH-associated macrophages (NAMs), as well as altered vascular signaling (Xiong *et al.*, 2019). It is also established that drugs and toxicants elicit spatial (zonal) toxicities such as acetaminophen which primarily affects the centrilobular region due to higher expression levels of xenobiotic metabolizing enzymes (Anundi *et al.*, 1993). Conversely, TCDD elicits periportal hepatotoxicity (Boverhof et al., 2006), despite preferential accumulation in the centrilobular region due to sequestration by induced CYP1A2 levels (Santostefano *et al.*, 1999). It remains unclear how transcriptional networks are implicated in these zonal toxicities. Single cell analysis is expected to further elucidate the role of specific cell (sub)populations in toxicity and models of liver disease progression.

Preclinical drug or chemical toxicity assessments typically involve dose-response designs which present numerous challenges for implemeting scSeq, primarily the required use of freshly isolated cells. Ongoing efforts to minimize experimental animal use also demand that studies maximize the extraction of relevant data including gross pathology, clinical chemistry, and histopathology. The impact of diurnal rhythm in large studies places additional logistic constraints regarding sample collection within a specific time window (Fader *et al.*, 2019). Most importantly, it is difficult to predict how treatment and/or disease pathologies impact cell populations, structure and viability, digestion efficacy, and cell type selection for single cell analysis without extensive *a priori* validation. Nuclei-based approaches address many of these challenges and produce results comparable to single cell approaches (Habib *et al.*, 2017; Habib *et al.*, 2016; Lake *et al.*, 2018; Zeng *et al.*, 2016). Notably, single nuclei RNAseq (snSeq) can be performed on frozen samples, providing the best solution for traditional toxicology assessments. In the presented study, snSeq was used to evaluate the hepatic effects of TCDD. snSeq analysis using frozen liver samples showed TCDD-elicited cell population shifts and cell-specific differential gene expression consistent with the progression of hepatic steatosis to NASH with fibrosis.

## MATERIALS AND METHODS

### Animals and treatment

Male C57BL/6 mice aged postnatal day (PND) 25 from Charles River Laboratories (Portage, MI) were housed in Innocages (Innovive, San Diego, CA) with ALPHA-dri bedding (Shepherd Specialty Papers, Chicago, IL) at 30-40% humidity and a 12h light/dark cycle. Animals were fed *ad libitum* Harlan Teklad 22/5 Rodent Diet 8940 (Harlan Teklad, Madison, WI), with free access to Aquavive water (Innovive). PND 28 mice were orally gavaged with sesame oil vehicle (Sigma-Aldrich, St. Louis, MO) or 30 μg/kg TCDD (AccuStandard, New Haven, CT) every 4 days for 28 days (7 treatments total). On day 28 (PND 52), animals were euthanized by CO2 asphixiation, and livers were immediately collected, frozen in liquid nitrogen, and stored at −80°C. All procedures were approved by the Michigan State University Institutional Animal Care and Use Committee.

### Nuclei isolation

Nuclei were isolated from frozen liver samples (~200mg) as previously described (dx.doi.org/10.17504/protocols.io.3fkgjkw). Briefly, livers were diced in EZ Lysis Buffer (Sigma-Aldrich), homogenized using a disposable dounce homogenizer, and incubated on ice for 5 minutes. The homogenate was filtered using a 70 μm cell strainer, transferred to microcentrifuge tube, and centrifuged at 500 x g and 4°C for 5 minutes. The supernatant was removed and fresh EZ lysis buffer was added for an additional 5 minutes on ice following by centrifugation at 500 x g and 4°C for 5 minutes. The nuclei pellet was washed twice in nuclei wash and resuspend buffer (1X PBS, 1% BSA, 0.2 U/μL RNAse inhibitor) with 5 minute incubations on ice. Following the washes, the nuclei pellet was resuspended in nuclei wash and resuspend buffer containing DAPI (10 μg/mL). The resuspended nuclei were filtered with 40 μm strainer and immediately underwent fluorescence activated cell sorting (FACS) using a BD FACSAria IIu (BD Biosciences, San Jose, CA) with 70 μm nozzle at the MSU Pharmacology and Toxicology Flow Cytometry Core (drugdiscovery.msu.edu/facilities/flow-cytometry-core). Sorted nuclei were immediately processed for snSeq.

### Single nuclei sequencing and data analysis

Libraries were prepared using the 10X Genomics Chromium Single Cell 3’ v3 kit (10X Genomics) and submitted for 150 bp paired-end sequencing at a depth ≥ 50,000 reads/cell using the HiSeq 4000 at Novogene (Beijing, China). Raw sequencing data were deposited in the Gene Expression Omnibus (GEO; Accession*ID TBD*). Following sequencing quality control, CellRanger v3.0.2 (10X Genomics) was used to align reads to a custom reference genome (mouse mm10 release 93 genome build) which included introns and exons to consider pre-mRNA and mature mRNA present in the nuclei.

Raw counts were further analysed using Seurat v3.1.1 (Butler *et al.*, 2018; Stuart *et al.*, 2019). Each sample was filtered for (i) genes expressed in at least 3 nuclei, (ii) nuclei which express at least 100 genes, and (iii) ≤1% mitochondrial genes. Additional quality control was performed using the scater package (v1.10.1). The DoubletFinder v2.0.2 package excluded putative doublets from subsequent analyses. Clustering of nuclei was performed using Seurat integration tools at a resolution of 0.2 and annotated using a semi-automated strategy by comparison to three published datasets [isolated hepatocytes sequenced by massively parallel single cell RNAseq (MARS-seq; GSE84498); isolated hepatic nonparenchymal cells (NPCs) by MARS-Seq (GSE108561); isolated NPCs using the 10X Genomics platform (GSE129516)]. Marker genes for individual nuclei clusters were also manually examined to verify annotation. Trajectory inference of hepatocytes was performed using Slingshot (Street *et al.*, 2018). Bulk RNAseq datasets from TCDD-treated mice were previously published (GSE109863 and GSE87543). Analysis code is available at github.com/zacharewskilab. The final snSeq Seurat object can be downloaded at Dataverse (dataverse.harvard.edu/naultran).

### Gene set enrichment analysis

Gene set enrichment analysis was performed on transcriptomes of individual nuclei using the Gene Set Variation Analysis (GSVA v1.30.0) package (Hanzelmann *et al.*, 2013) with a minimum of 10 genes and maximum of 300 gene per gene set. Mouse gene sets were obtained from the Gene Set Knowledgebase (GSKB; ge-lab.org/gskb/) filtered to only include MSIGDB, GENESIGDB, SMPDB, GO, KEGG, REACTOME, EHMN, MIRNA, MICROCOSM, MIRTARBASE, MPO, PID, PANTHER, BIOCARTA, INOH, NETPATH, WIKIPATHWAYS, MOUSECYC, TF, and TFACTS gene sets (6,513 total). Human-based (e.g. HPO), exposure-based (e.g. DRUGBANK), or gene sets with limited annotation (e.g. LIT) were excluded. Uniform Manifold Approximation and Projection (UMAP) reduction using the first 30 principal components analysis (PCA) dimensions was used for visualization using original cell type annotations. For the visualization of marker and differentially enriched pathways, a network of gene sets was generated using the Enrichment Map plugin for Cytoscape (Merico *et al.*, 2010). Nodes were summarized using AutoAnnotate (Kucera *et al.*, 2016) and subsequently manually curated. Nodes were redrawn using the Enhanced Graphics package.

## RESULTS

### snSeq identifies major liver cell (sub)types in vehicle and TCDD treated mice

We adapted a single cell RNAseq protocol for frozen cancer tissue biopsy samples that was compatible with the 10x Genomics technology to characterize gene expression in nuclei isolated from mouse liver samples treated with either sesame oil vehicle or 30 μg/kg TCDD. A total of 16,015 individual nuclei transcriptomes (9,981 and 6,034 in vehicle and TCDD treated, respectively) were characterized across 2 biological replicates per treatment group after quality control and doublet removal. The average number of expressed genes detected in each sample was 17,920, consistent with our published bulk RNAseq assessments collected using a similar study design (Fader *et al.*, 2017a; Nault *et al.*, 2015). The median number of unique detected genes in individual nuclei was 1,694 with a median unique molecular identifier (UMI; transcript) count of 3,385 per nuclei. As expected with a nuclear preparation, there was negligible expression of mitochondrial genes. Long non-coding RNAs (lncRNAs) such as *Malat1* and *Gm42416* were most abundantly expressed as reported in other scSeq and snSeq studies (Habib *et al.*, 2017; Zeng *et al.*, 2016). The number of unique expressed genes and median UMI count (1,500 and 3,805, respectively), as well as total expressed genes (19,907), was similar to a published 10x Genomics liver single cell dataset (19,349) (Xiong *et al.* (2019), lending further confidence in our snSeq approach. CellRanger detected some ambient RNA contamination in some samples, likely due to the lysis of cells during nuclei isolation (Alvarez *et al.*, 2019). However, neither of the common indicators of ambient RNA contamination (mitochondrial or hemoglobin gene expression) were elevated in our samples. Consequently, we did not use any ambient RNA decomtamination tools such as DecontX, SoupX or DIEM (Alvarez *et al.*, 2019; Yang *et al.*, 2019; Young and Behjati, 2020).

Integration and clustering of nuclei transcriptomic profiles identified eleven clusters, of which only the neutrophil cluster was unique to TCDD treatment (**Fig. 1A, D**). Nuclei clusters were initially annotated by integrating our dataset with two published single cell RNA sequencing datasets and comparing annotation assignments (Halpern *et al.*, 2018; Xiong *et al.*, 2019) (**Fig. 1B**), and subsequently corroborated using the expression of known markers for specific cell types based on published reports and/or panglaoDB (*Franzén et al., 2019*). For example, nuclei with higher levels of Stabilin 2 (*Stab2*) (**Fig. 1B-C**), an expression marker for hepatic sinusoidal endothelial cells (Nonaka *et al.*, 2007), were identified as “Endothelial Cell” nuclei in our dataset, as reported by Xiong *et al.* (2019) in a diet induced NASH model. Similarly, cholangiocyte expression profiles aligned with a distinct nuclei cluster (**Fig. 1B**) (Xiong *et al.* (2019). Although the classical cholangiocyte marker, SRY-Box 9 (*Sox9*), was not expressed in our nuclei, the similarity between the expression profiles in our dataset and cells expressing *Sox9* reported by Xiong *et al.* (2019) suggested that these nuclei are indeed cholangiocytes. Examination of distinguishing markers for each nuclei cluster (**Fig. 1C**) indentified *Pkhd1* (also known as fibrocystin) as a marker for cholangiocytes. *Pkhd1* has been shown to be expressed in rat cholangiocytes (Masyuk *et al.*, 2003). Other distunguishing markers such as *Ebf1*, important in B cell development (Zhang *et al.*, 2003), and the T cell adaptor *Skap1* (Smith *et al.*, 2016), were in agreement with cluster annotations.

**Figure 1.**
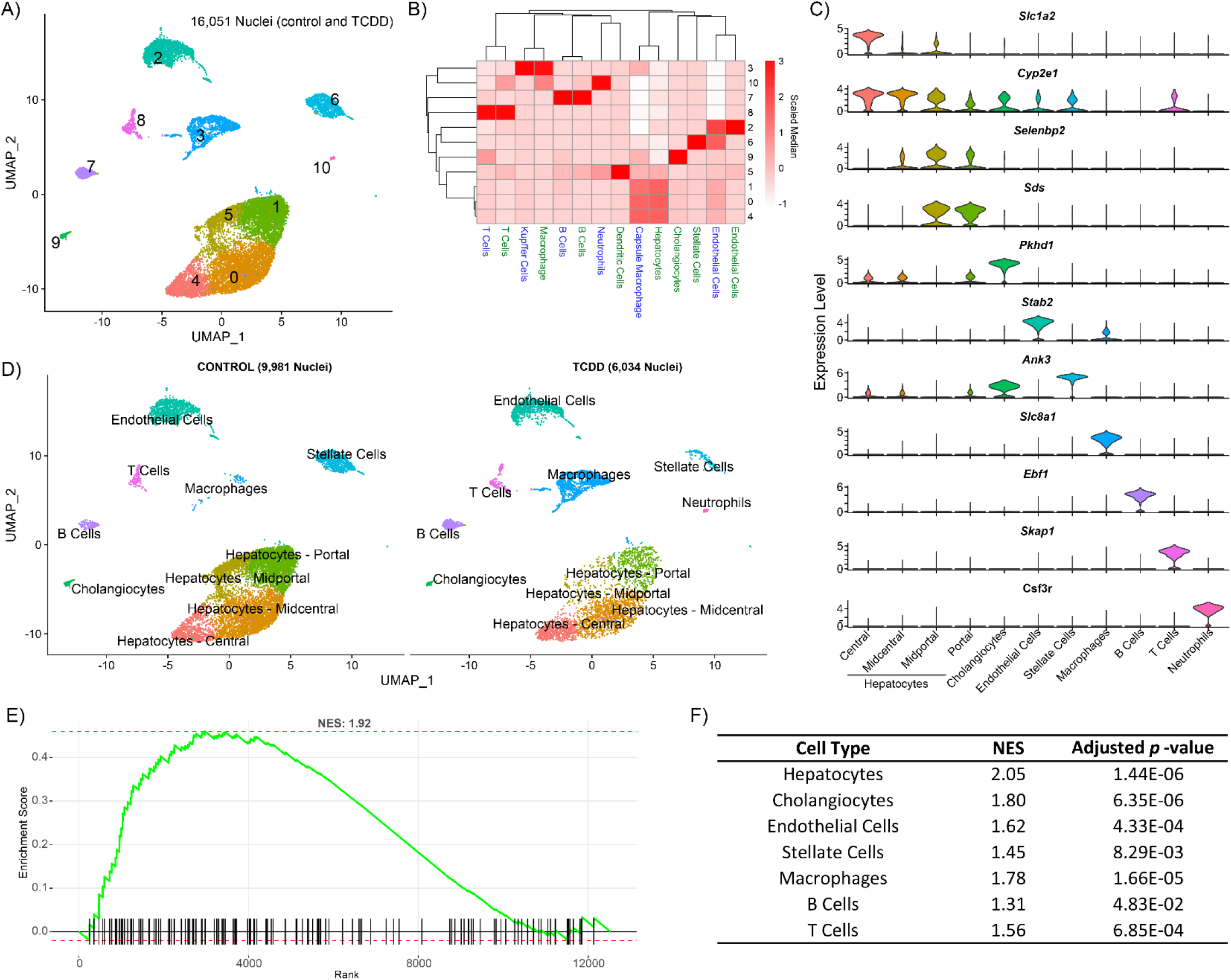
Single nuclei RNAseq (snSeq) analysis of nuclei isolated from frozen liver samples of mice gavaged every 4 days for 28 days with sesame oil vehicle or 30 μg/kg TCDD. (A) Uniform Manifold Approximation and Projection (UMAP) visualization of nuclei isolated from vehicle and TCDD-treated liver samples clustered based on gene expresion profile similarity. (B) Label transfer prediction from published liver single cell RNA seq data (Halpern et al. (2018) in blue, Xiong et al. (2019) in green) was used to identify similar clusters for annotation. (C) Expression distribution for distinguishing (largest average fold-change, adjusted *p*-value ≤ 0.05) marker genes within each specific cell type. Color of clusters in (C) correspond to cell type clusters in (A). (D) UMAP visualization of annotated nuclei in control and TCDD treated samples. Gene set enrichment analysis (GSEA) of nuclear biased genes (1^st^ quartile) (Bahar Halpern *et al.*, 2015) using genes ranked from most to least abundantly expressed in our nuclei dataset compared to Xiong et al. (2019) whole cell data was performed for (E) all cell types and (F) paired cell types identified in both datasets. NES represents the normalized enrichment score.

Our dataset suggests *Sox9* may be less abundant in nuclei sample preparations, consistent with previous reports indicating some genes exhibit nuclear or cytosolic biases (Bahar Halpern *et al.*, 2015; Habib *et al.*, 2017; Zeng *et al.*, 2016). Comparison of our dataset to Xiong *et al.* (2019) using a similar model and the same technology shows excellent concordance in identifying genes expressed in liver cells (16,789; 84 - 87%) despite Xiong et al.’s enrichment for non-parenchymal cells. However, it is evident that relative levels of certain genes differ, particularly lncRNA’s. Comparisons of nuclear and cytoplasmic gene expression in liver cells (mixed cell types) identified *Mlxipl* (ChREBP) and *Nrlp6* as retained nuclear genes, as well as the lncRNA *Neat1* (Bahar Halpern *et al.*, 2015). In agreement, our dataset exhibited greater expression of these genes compared to whole cell liver expression data (Xiong *et al.* (2019) (**Fig. S1**). We performed GSEA on ranked genes from nuclear biased (elevated fold change in our vehicle control dataset compared to control whole cell data) to cytosolic/whole-cell biased (elevated in control single cell data compared to our vehicle control dataset) using nuclear biased genes (top 1^st^ percentile) identified by Bahar Halpern *et al.* (2015) as the gene set. **Fig. 1E-F** demonstrates our single nuclei gene expression dataset is indeed enriched in nuclear biased genes in all cell types.

### TCDD shifted cell type proportions

TCDD elicited shifts in the relative proportions of nuclei clusters (**Fig. 2A**). Macrophages increased from 0.5% in controls to 24.7% in TCDD treated samples, accompanied by other immune cells such as B cells (1.4% to 7.5%), T cells (1.7% to 4.7%), and neutrophils (0.0% to 1.4%). This is consistent with TCDD-induced inflammation (**Fig. 2B**) and elevated levels of proinflammatory cytokines, such as TNFα and IL-6 (Nault *et al.*, 2016). Overall, the relative proportion of hepatocytes in TCDD treated samples was reduced by 46.6% (75.3% in controls compared to 40.2% in treated samples). All zones were reduced except for central hepatocytes which increased from 8.6% to 10.8% (**Fig. 2A**). It should be noted that distinguishing whether decreases in relative proportion of cell types reflect cell death, loss of cell types during processing, and/or a true shift in relative levels is challenging, although previous studies using the same study design did not report cell death from histological evaluation (Fader *et al.*, 2017b). These challenges are further explored in the discussion below.

**Figure 2.**
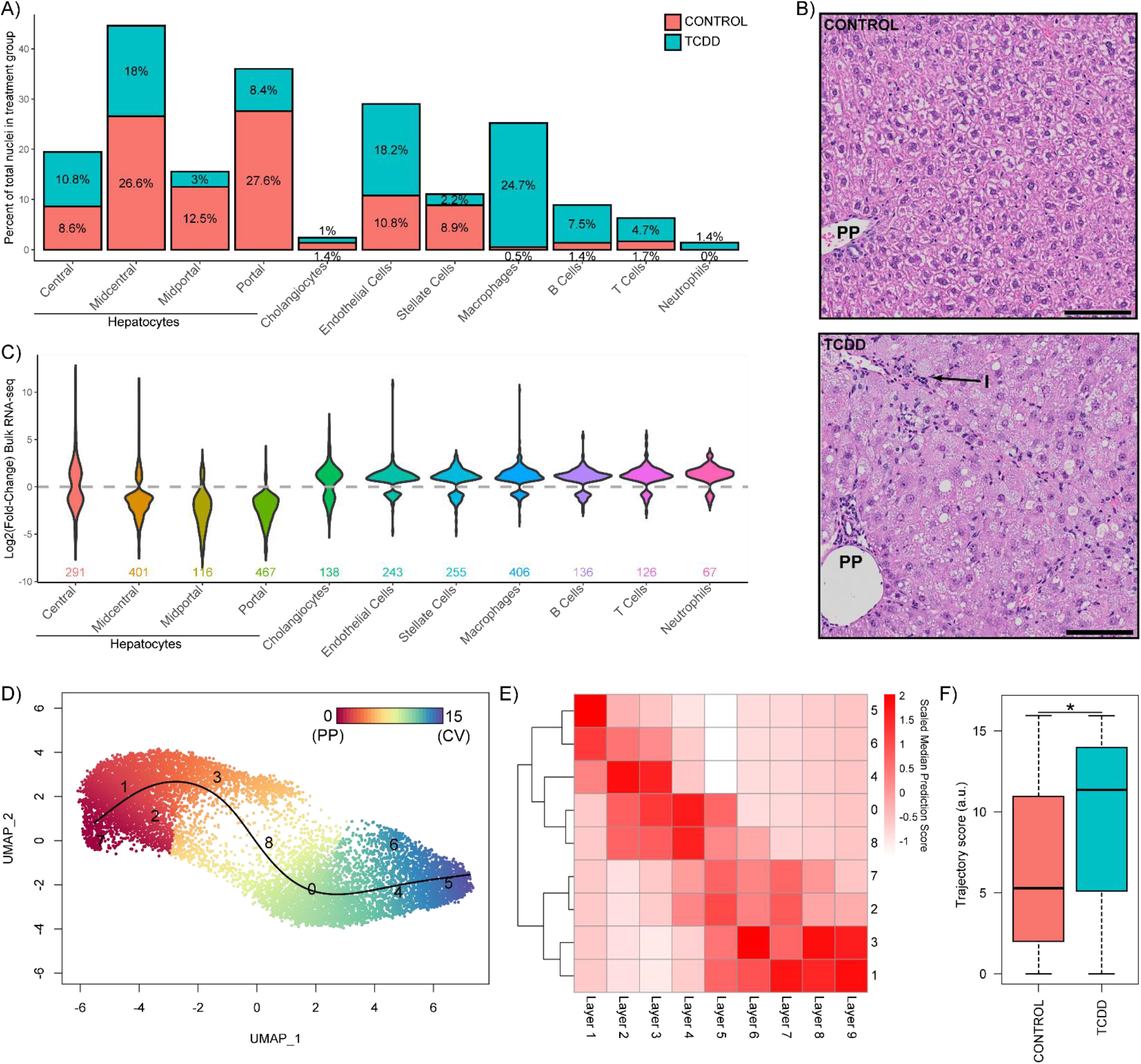
Hepatic cell population shifts in response to TCDD. (A) Percentage of nuclei represented in cell-type specific clusters in control and treated samples. (B) Representative liver photomicrographs from control and TCDD treated samples using the same study design (Fader et al., 2017a) showing periportal (PP) immune cell infiltration (I). Scale bar represents 100 μm. (C) Violin plots of fold-change expression distribution for genes defining cell type clusters (adjusted p-value ≤ 0.05) in TCDD-elicited bulk RNAseq dataset. (D) Reclustering and trajectory analysis of only hepatocyte nuclei from vehcile control and TCDD treatment groups using a resolution of 1.2 to identify 9 clusters in order to (E) compare snSeq gene expression to the 9 distinct spatially resolved hepatocyte layers defined by Halpern et al. (2017). (F) Nuclei distribution along the trajectory determined in control and TCDD treated samples. The asterisk (*) indicates a significant difference in distribution with TCDD causing a shift in the hepatocyte nuclei population towards the central cluster/layer as determined by the Kolmogorov-Smirnov test (*p* ≤ 0.05).

To further investigate cell population shifts, we re-examined our published hepatic bulk RNAseq (same species, strain, sex, age, treatment regimen, dose, and duration of exposure; (Fader *et al.*, 2017a)) for TCDD-elicited fold-change distribution of genes identified as cell type markers (adjusted p-value ≤ 0.05) (**Fig. 2C**). Overall, marker gene expression for midcentral, midportal, and portal hepatocytes was repressed in our bulk RNAseq dataset as evidenced by the bulk RNA-seq log2(fold-change) distribution ≤ 0 (wider below the gray line). Conversely, marker genes for cholangiocytes, endothelialcells, stellate cells, and immune cells were induced. Comparing the changes in relative proportions (**Fig. 2A**) to differential gene expression in bulk RNAseq (**Fig. 2C**) confirms the impact of TCDD on cell population shifts in bulk RNAseq analysis. Specifically, central hepatocyte marker genes were repressed despite a modest increase in relative cell numbers suggesting TCDD effects on gene expression. In contrast, macrophage marker induction concommitant with increased macrophage number makes it difficult to distinguish a direct effect on gene expression from an increase in the number of infiltrating cells with basal gene expression. The limited ability to delineate these effects using bulk RNAseq demonstrates another advantage of a single cell/nuclei analysis.

### Characterization of spatially resolved hepatocytes

To investigate zonal gene expression using snSeq data, hepatocyte nuclei from both treatment groups were selected, re-integrated, and re-clustered to obtain 9 clusters, guided by the 9 hepatocyte zones (*i.e.* layers) defined by Halpern *et al.* (2017) (**Fig. 2D**). Our hepatocyte nuclei clusters demonstrate high similarity to the transcriptomes of the different cell layers defined by Halpern *et al.* (2017) (**Fig. 2E**). Specifically, nuclei cluster labeled “5” was most similar to layer 1 cells while clusters “1” and “3” were most like layers 8 and 9. Zonal trajectory determined using Slingshot (Street *et al.*, 2018) shows a transition in gene expression profiles from cluster “5” to “1”, reflecting a gradient from central (layer 1) to portal (layer 9) (**Fig. 2D-E**). Cluster “7” was less defined, though closer examination identified the expression of portal markers, along with elevated expression of metallothioneins (*Mt1* and *Mt2*) equally expressed in control and treated samples (**Fig. S2**). Comparing zonal distribution between control and treated nuclei revealed a shift towards central hepatocyte expression, suggesting TCDD caused a loss of portal hepatocyte gene expression, or portal hepatocyte were lost during sample preparation (**Fig. 2F**). TCDD elicits periportal hepatotoxicity, and therefore the loss of identity from de- or trans-differentation cannot be distinguished from cell-specific cytotoxicity.

### Differential gene expression elicited by TCDD is largely cell type specific

Of the 19,907 genes detected across all liver cell-types, 10,951 were identified as differentially expressed, comparable to the 9,313 (of 19,935) differentially expressed genes reported in our comparable bulk RNAseq study (Fader *et al.*, 2017a). Hepatocyte sub-types exhibited the greatest similarity in differential gene expression with 92.6% (1,305/1,409) of midportal differentially expressed genes (DEGs) overlapping with portal DEGs (**Fig. 3A**). However, this only represents 20.6% (1,306/6,346) of portal DEGs suggesting midportal cells may be a subset of portal hepatocytes. Cholangiocytes compared to macrophages shared the fewest overlapping DEGs with only 38 in common (31.1% and 11.8% of DEGs, respectively). Only 28 genes were differentially expressed in all cell types, with 26 repressed in every cell type. Only Cadherin 18 (*Cdh18*) was induced in all cells, while Dihydropyrimidine dehedrogenase (*Dpyd*) was induced in hepatocytes but repressed in all other cell types (**Fig. 4A**).

**Figure 3.**
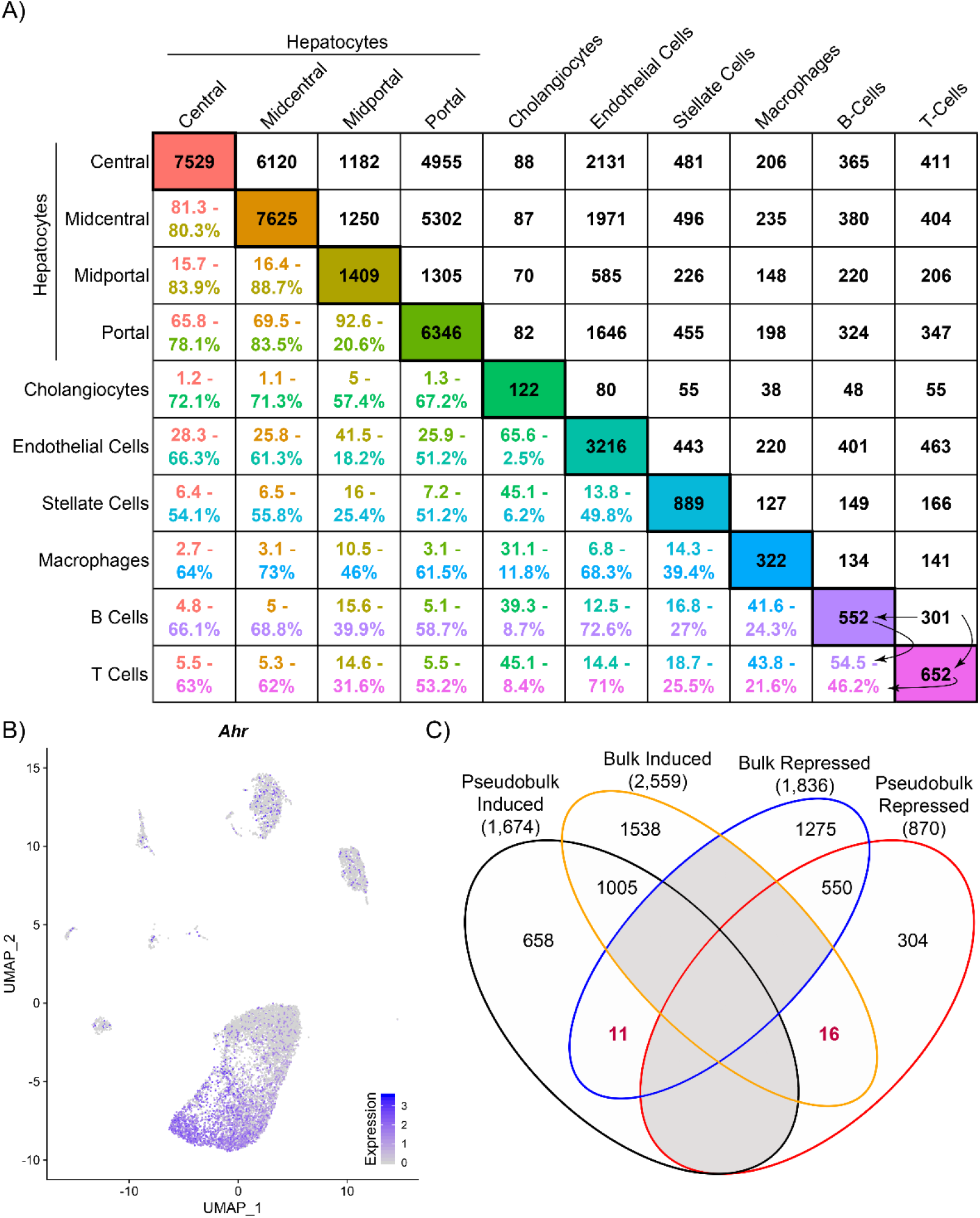
TCDD-elicited differential gene expression in distinct hepatic cell sub-types. (A) Cross-comparisons of differentially expressed genes (DEGs; adjusted p-value ≤) between cell (sub)types. The number of DEGs (adjusted *p*-value ≤ 0.05) determined for each nuclei cluster is provided in the diagonal boxes color-coded according to cell sub-types identified in **Fig. 1A**. The number of common DEGs determined between each pair of cell types is provided in upper right portion of table in black font. The percentage of common DEGs between cell types is provided in the bottom left portion of table using two different colored fonts. The upper colored font represents the percent overlap of DEGs relative to the cell-subtype for the column while the lower colored font represents the percent overlap of DEGs relative to the cell-subtype for the row. An example of percent calculations is shown between B cells and T cells using arrows (e.g. 301/552 = 0.545). (B) UMAP visualization of *Ahr* expression in all nuclei from vehicle control samples. (C) Comparison of pseudobulk and bulk RNAseq analyses for up-regulated and down-regulated genes detected in both datasets.

**Figure 4.**
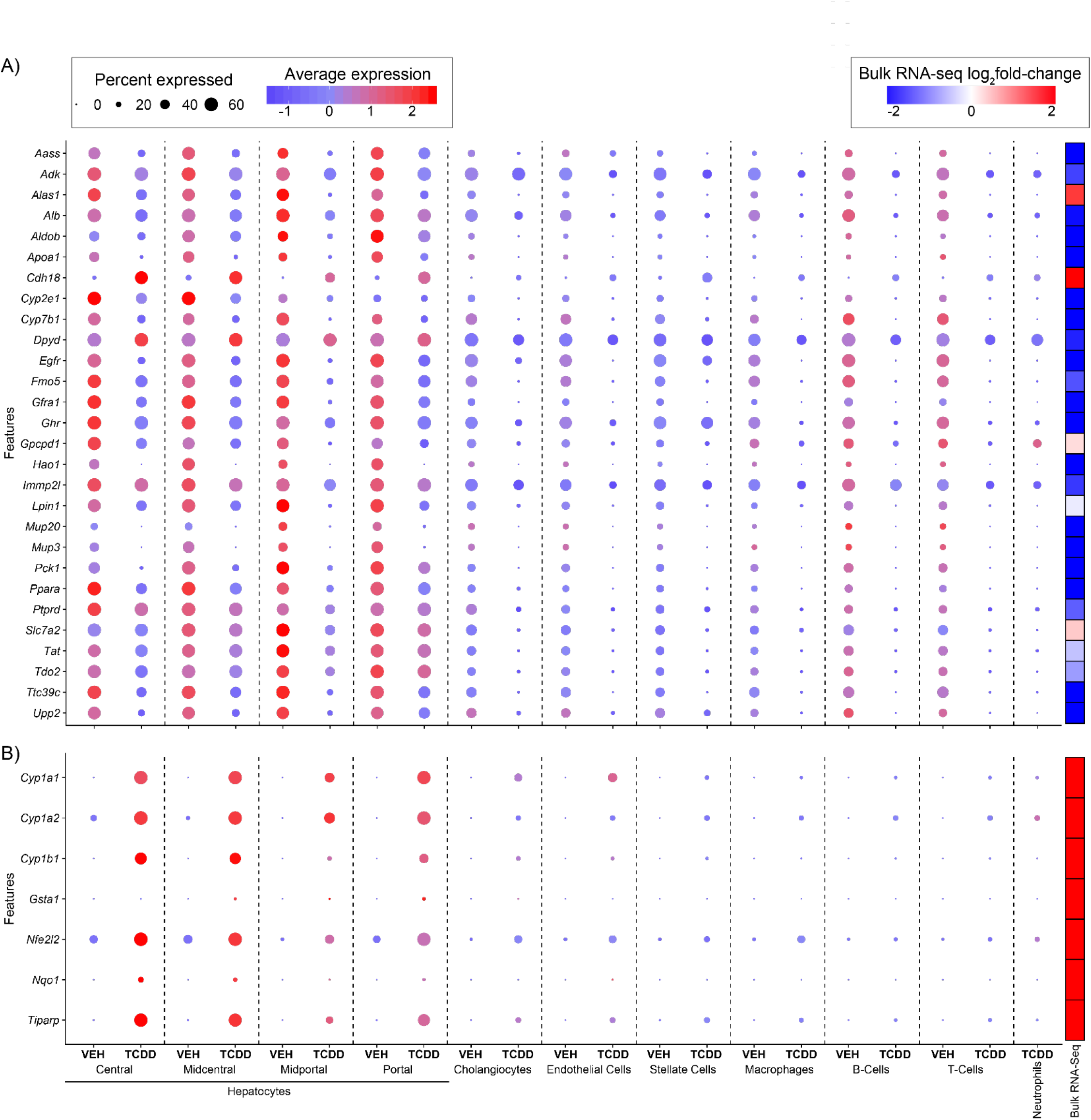
DEGs across cell (sub)types. Relative expression and percent nuclei expressing (A) differentially expressed genes across all cell types and (B) AhR battery genes. Dot size represents the % of nuclei expressing DEG. Color scale (red to blue) of dots reflect average expression level. Last column on right indicates the gene fold change in the bulk RNAseq dataset at 30 ug/kg TCDD collected using the same model and study design. Neutrophils are not shown as they were only dected in TCDD treated samples.

The number of DEGs in hepatocytes correlated with basal *Ahr* expression levels in control samples. Central and midcentral hepatocytes exhibited the highest basal *Ahr* expression levels (detected in 79% and 65% of nuclei, respectively (**Fig. 3B**)) and exhibited the most DEGs. Portal hepatocytes also exhibited a large number of DEGs (6,346) despite a lower AhR expression level (detected in only 17% of portal nuclei) suggesting secondary or tertiary factors (e.g. chromatin accessibility, metabolite gradient, cell-cell interactions) contributed to TCDD responsiveness. Induction of classic AhR target genes was not detected in all cell types (**Fig. 4B**) and exhibited fold-change differences between cell types. For instance, *Cyp1a1* was induced 2,736-fold in portal hepatocytes, consistent with reports in bulk RNAseq analyses. The 1,177-fold repression of *Ces3b* reported in bulk RNAseq was comparable to 3,089-fold repression observed in central hepatocytes. Many genes went from undetected to detected, confounding fold-change estimates (*i.e.*, division by zero), though these genes were typically significantly induced in bulk RNAseq. For example, *Gpnmb* was induced 1,206-fold in our bulk RNAseq dataset but only detected in macrophage following TCDD treatment in the snSeq dataset. A large number of zeroes is expected in single cell/nuclei RNAseq resulting from either absent expression and/or dropouts.

To compare single nuclei to bulk RNAseq data, snSeq data was first converted to “pseudo-bulk” data by summing read counts across all nuclei to approximate bulk expression. Differential gene expression analysis for our pseudo-bulk data identified 2,812 differentially expressed genes (DEGs; adjusted *p*-value ≤ 0.05 & |fold change| ≥ 2), 2,544 of which were also detected in bulk RNAseq. Of which 1,005 were induced in both analyses with 550 repressed (**Fig. 3C**). Only 27 genes showed divergent expression (e.g. induced in one dataset and repressed in the other). Due to the substantial technological differences in determining transcript counts, some differences are not surprising. In addition to technical differences, some genes may differ due to diurnal rhythm (e.g. *Alas1*) or cell type biases (e.g. macrophage genes *Adgre4* and *Clec4f*).

### TCDD elicits unique functional changes in cell populations

To identify enriched cellular pathways, processes, and functions associated within specific cell (sub)types, all constitutively and differentially expressed genes were analyzed in control and treated samples. Enrichment scores were calculated for 6,513 gene sets obtained from the Gene Set Knowledgebase (GSKB) for each nucleus independent of cell type classification. UMAP visualization of functional enrichment higlights TCDD-elicited shifts in enriched functions within specific cell types, most notably hepatocytes (**Fig. S3**). Pathways, processes, and functions enriched within specific cell (sub)types were consistent with their known physiological roles (**Fig. 5; inner ring**). For instance, macrophages were enriched in phagocytosis and inflammation gene sets while endothelial cells were enriched in vasculature related genes. Hepatocytes were largely enriched in genes associated with the metabolism of carbohydrates, lipids, amino acids, and bile acids, as well as one carbon metabolism, and were consistent with their expected zonal distribution. Specifically, amino acid metabolism and the coagulation cascade were enriched in the portal region while lipid, bile acid, and phase I xenobiotic metabolism were centrally enriched (**Fig. 5**), in agreement with recent zonal proteomics results (Ben-Moshe *et al.*, 2019).

**Figure 5.**
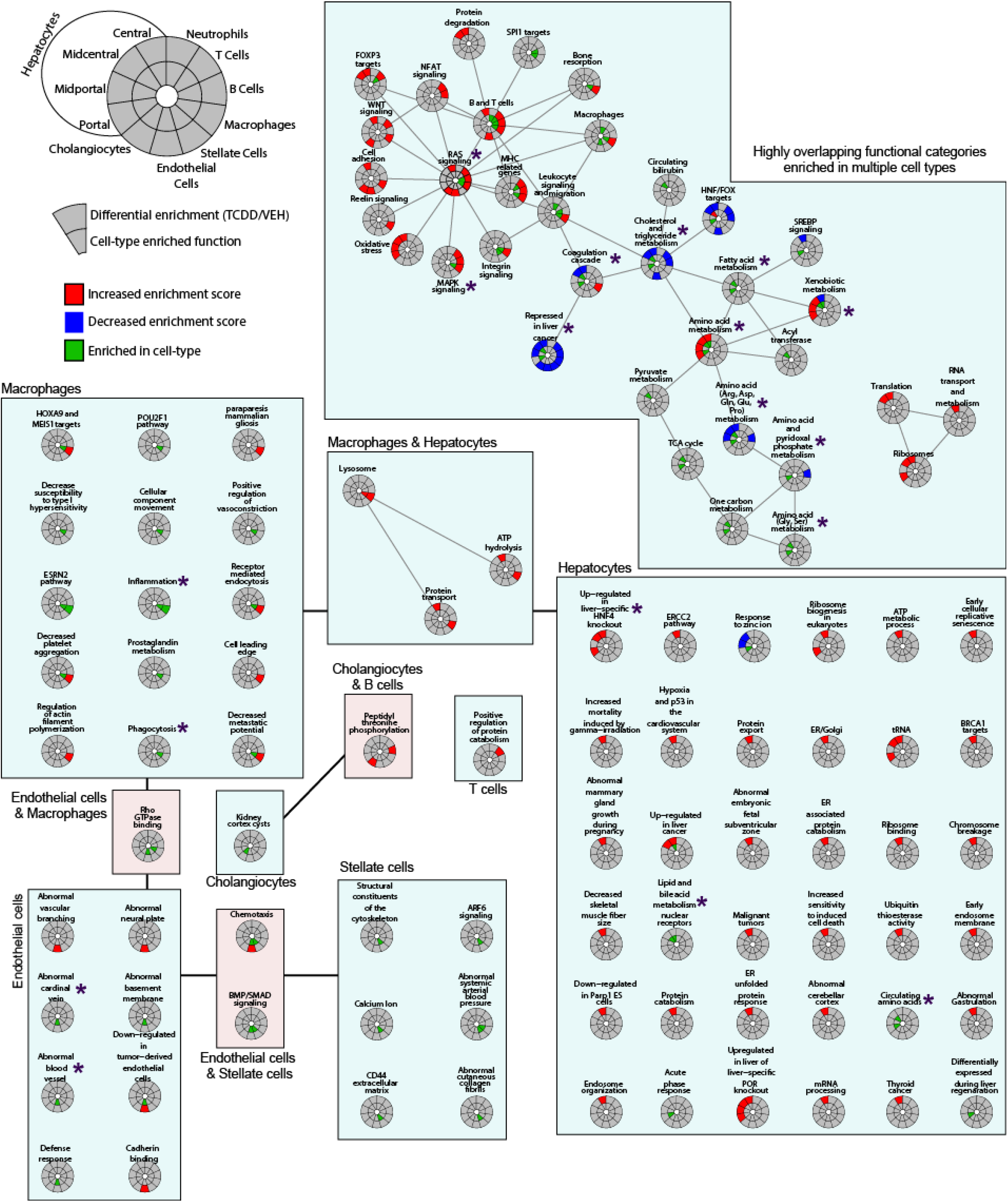
Gene set enrichment analysis of single-nuclei transcriptomes isolated from frozen liver samples of mice gavaged every 4 days for 28 days with sesame oil vehicle or 30 μg/kg TCDD. Circos plots of functional enrichment analysis for each cell type indicates marker functions (adjusted *p*-value ≤ 0.05 compared to other cell types; inner ring; green) and functions showing increased (red) or decreased (blue) enrichment scores following TCDD treatment (outer ring). Enrichment scores for 6,513 gene sets were determined using GSVA, grouped based on gene membership similarity using Enrichment Map, and summarised using AutoAnnotate followed by manual curation as described in the materials and methods. Asterisks (*) indicate functions discussed in the manuscript.

We next examined nuclei for functional enrichment following TCDD treatment using enrichment score differences (fold-change) between control and treated nuclei (**Fig. 5; outer ring**). RAS signaling was the most highly connected nodes suggesting high similarity in gene membership with other nodes, particularly those associated with nonparenchymal cells. Metabolism pathways were largely associated with hepatocytes, including repression of cholesterol and triglyceride metabolism, and metabolism of selected amino acids which were either induced or repressed (e.g. Arg, Asp, Gln, Glu, and Pro). Not surprisingly, xenobiotic metabolism was induced in all hepatocytes except central hepatocytes. This is due to the repression of several cytochrome P450s and high constitutive expression of phase I metabolism genes in central hepatocytes. Other notable gene sets induced in hepatocytes were i) “up-regulated in liver-specific HNF4α knockout” suggesting TCDD repressed HNF4 signaling, and ii) “repressed in liver cancer” suggesting transcriptomic changes consistent with hepatocarcinogenicity of TCDD.

### Emergence of new macrophage (sub)types following treatment

A hallmark of TCDD exposure is the infiltration of immune cells into the liver (Fader *et al.*, 2017b). **Fig. 1D** and **2B** show the increased presence of macrophages and neutrophils following treatment with TCDD. Resident macrophages in a normal liver largely consist of Kupffer cells (KCs) marked by high *Adgre1* (F4/80) expression. Upon injury, motile monocyte derived macrophages (MDMs) with high levels of *Itgam* (Cd11b) and *Ccr2* expression, are recruited (Dong *et al.*, 2019; Xiong *et al.*, 2019). Further analysis of macrophage nuclei identified five distinct clusters, of which four expressed high levels of *Adgre1* and *Cd5l* (**Fig. 6**) while none exhibited high levels of *Itgam* (not detected) or *Ccr2* (expressed in ≤ 4% of control macrophages). Macrophage clusters 1, 2, and 3 not only expressed high levels of *Adgre1* and *Cd5l* but also *Gpnmb* which was not present in KCs from control samples. Interestingly, comparable macrophage clusters were reported in a diet-induced NASH mouse model, described as NASH associated macrophages (NAMs), that also exhibited high *Trem2* expression (Xiong *et al.*, 2019). Although highly induced in our bulk RNAseq dataset, we did not identify a macrophage population expressing high *Trem2* levels, nor *Cd9*, as reported in NAMs by Xiong *et al.* (2019), and is likely another example of a gene biased to the cytoplasm. Indeed, both genes are more abundant in the cytosol than nuclei in liver cells (Bahar Halpern *et al.*, 2015). KC cluster 3 was also determined to express high levels of the cell cycle gene *Top2a*, possibly identifying a proliferating NAM subpopulation. The fourth and smallest cluster expressed high levels of *Bcl11a* and *Ccr9* (**Fig 6B**), and likely represent plasmacytoid dendritic cells as previously reported (Halpern *et al.*, 2018).

**Figure 6.**
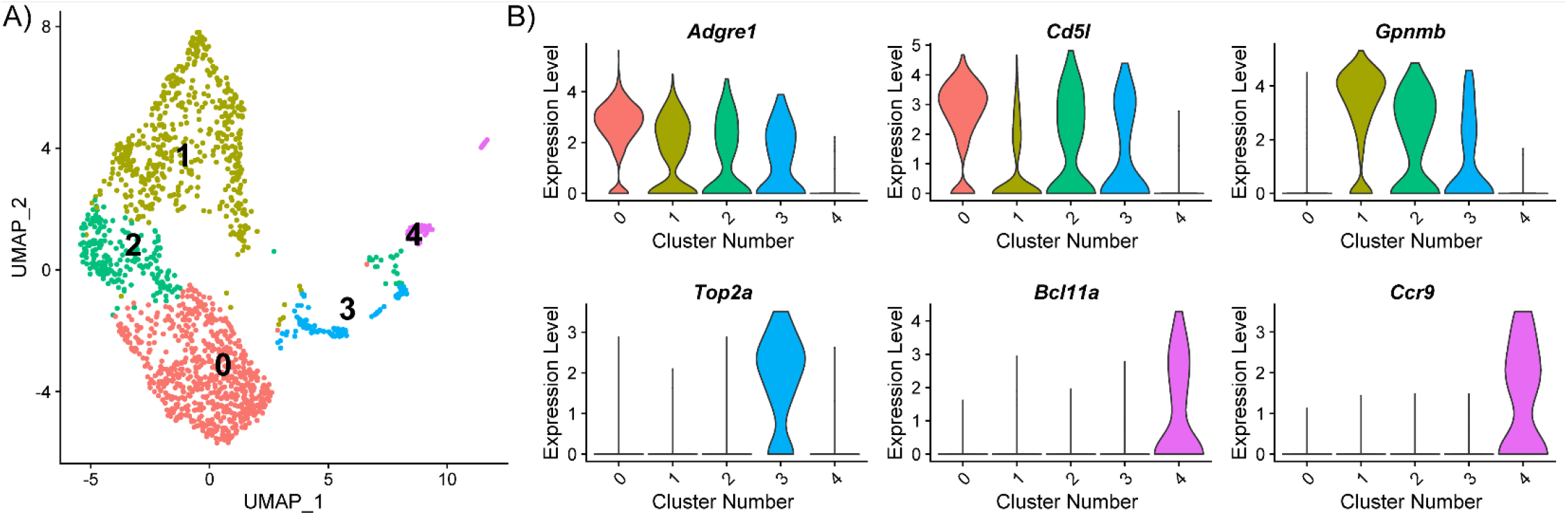
Clustering of macrophage nuclei isolated from frozen liver samples of mice gavaged every 4 days for 28 days with sesame oil vehicle or 30 μg/kg TCDD. (A) UMAP plot of macrophage nuclei determined at 0.2 resolution using Seurat. (B) Violin plots of the expression levels of unique genes that distinguish macrophage subtypes.

## Discussion

Single cell/nuclei transcriptomics has not only provided novel insight into molecular mechanisms involved in cellular development and normal cell function, but has also identified and characterized rare cell (sub)types (Habib *et al.*, 2016; Halpern *et al.*, 2018; Halpern *et al.*, 2017; Iacono *et al.*, 2019; Su *et al.*, 2017; *Xiong et al., 2019*). It has the potential to further elucidate the mechanisms of toxicity elicited by exogenous agents that disrupt normal cellular functions. Most importantly, it enables the assessment of cell type-specific responses, thus integrating systemic (multi-organ) and local events within the context of a tissue or organ. In this study, we used a nuclei-based strategy to minimize potential biases in cell (sub)types due to toxicity that was also logistically feasible within a large dose-response and time-course study. Similar strategies have been used for neurons which vary wildly in shape, density, and other cellular characteristics, as well as *post-mortem* human tissues with good success (Bakken *et al.*, 2018; Krishnaswami *et al.*, 2016).

Single cell analyses have been performed on mouse and human liver samples, though most have enriched for either hepatocytes or non-parenchymal cells (NPCs) (Halpern *et al.*, 2018; Halpern *et al.*, 2017; MacParland *et al.*, 2018; Xiong *et al.*, 2019). Using a single nuclei approach, we captured the expected hepatic cell types identified in previous studies with hepatocyte nuclei highly represented, given they compose ≥ 80% of the liver cell population. Consequently, immune cells, particularly in control samples, were difficult to identify, such as the plasmacytoid dendritic cells and capsule macrophages identified in NPC enriched samples (Halpern *et al.*, 2018). A potential solution to counter this unequal hepatocyte sampling bias could be FACS separation of nuclei based on DNA, as NPCs are enriched in the diploid (2N) population while hepatocyte nuclei are often polyploid (Digernes and Bolund, 1979). However, TCDD alters liver cell ploidy (Moreno-Marin *et al.*, 2018), which may introduce biases in the relative proportions of distinct cell types, and therefore we did not separate nuclei by ploidy in this study. While our snSeq dataset could not estimate the total number of each cell type, the emergence of new cell populations and the transcriptomic changes of resident cells could be distinguished. Conversely, distinguishing loss of cell (sub)types due to toxicity or de-differentiation represents a unique challenge.

Our single nuclei strategy highlighted several advantages over single cell analysis to assess hepatotoxicants. First, nuclei isolated from frozen samples could cleary be classified as distinct known hepatic cell types. Second, despite higher representation of nuclear-biased genes, TCDD-elicited differential expression determined by bulk RNAseq was largely recapitulated in a pseudo-bulk version of our snSeq dataset with only 38 genes showing conflicting results, which could be due to other confounding factors (e.g. different technologies, diurnal rhythm). However, our dataset was enriched with nuclear biased genes while other cell type markers, such as the cholangiocyte marker *Sox9*, were not detected. Single nuclei droplets may also be more susceptible to ambient RNA contamination, though our mitochondrial gene expression content was low. Despite differences in mRNA capture, sequencing, counting, and analysis, the strong correspondence indicates snSeq is a viable alternative with distinct advantages over scSeq analysis.

Hepatotoxicants can elicit zonal toxicity based on their mode of action and the functional gradient of the lobule chords (central vein to portal vein). Toxicants not requiring bioactivation commonly damage periportal regions when first encountered via the portal vein. Conversely, toxicants requiring bioactivation are more often toxic to pericentral regions due to higher basal expression of phase I metabolism enzymes (e.g. cytochrome P450s). TCDD is a periportal hepatotoxicant, though damage spans the periportal to pericentral regions (panacinar) with increasing dose. Interestingly, pericentral hepatocytes were most responsive to TCDD, followed by periportal hepatocytes. TCDD primarily accumulates in central hepatocytes due to the induction of *Cyp1a2*, a phase I metabolism enzyme which sequesters TCDD (Santostefano *et al.*, 1999). The high dose of TCDD that caused panacinar damage together with the higher basal expression of *Ahr* in central hepatocytes likely coalesced to elicit the large number of differentially expressed genes in this region. TCDD also reduced the portal hepatocyte representation. Spatial transcriptomics approach can be used to further investigate the loss of periportal cells in future studies.

SnSeq also provided novel insight into cell-specific gene expression and the roles of distinct cell populations. RAS/MAPK signaling was activated in hepatic NPCs in response to TCDD to promote cell proliferation and cell survival (Yang and Liu, 2017). Previous studies have linked RAS/MAPK signaling to inflammation in human NAFLD, likely through the production of growth factors, cytokines, chemokines, and adhesion molecules (Jimenez-Castro *et al.*, 2019). Moreover, TCDD represses hepatocyte proliferation (Jackson *et al.*, 2014), and RAS induction by TCDD in HepG2 cells does not activate the downstream effector, MAPK (Yamaguchi and Hankinson, 2018). The lack of hepatocyte enrichment is also consistent with the inhibition of hepatocyte proliferation and infiltration of immune cells. Further investigation of macrophage subpopulations identified a high *Gpnmb*-expressing Kupffer cell population following TCDD treatment which was also reported in a diet-induced NASH model (Xiong *et al.*, 2019) and in humans with steatosis or NASH (Katayama *et al.*, 2015), revealing conserved molecular events across disparate etiologies and species. In agreement with the loss of liver-specific gene expression (Nault *et al.*, 2017), we find enriched genes that were also up-regulated in a liver-specific HNF4 knockout model, a transcription factor essential for liver-specific gene expression (Ohguchi *et al.*, 2008). Similarly, there was decreased expression of genes repressed in liver cancer, consistent with TCDD being a known mouse liver carcinogen. Decreased enrichment was observed in most cell types as the gene set was produced from whole tissue expression profiling, highlighting some caveats when examining functional changes using single cell/nuclei data.

In conclusion, snSeq represents a valuable strategy to characterize cell-specific responses of hepatotoxicants. Despite known biases in nuclear and cytosolic transcript levels, single nuclei transcriptomic data accurately characterize differential expression elicited by TCDD as reflected in (1) the consistency with bulk gene expression data, (2) the identification of NASH associated macrophages, and (3) changes in functional enrichment of pathways associated with TCDD-elicited hepatotoxicity. Furthermore, the analysis of frozen samples enables comprehensive dose- and time-dependent investigation of cell-specific adverse effects.

## Acknowledgements

The authors would like to thank Dr. Luciano Martelotto (University of Melbourne) for his help in optimizing the nuclei isolation protocol for single nuclei RNA sequencing, as well as the labs of Drs. Jiande Lin (University of Michigan) and Shalev Itzkovitz (Weizmann Institute of Science) for sharing processed datasets used to semi-automate cluster annotation. This work was supported by the National Institute of Environmental Health Sciences Superfund Research Program [NIEHS SRP P42ES04911] to TRZ and the National Human Genome Research Institute [NHGRI R21HG010789] to TRZ and SB. TRZ and SB are partially supported by AgBioResearch at Michigan State University.

---

**Figure S1.**
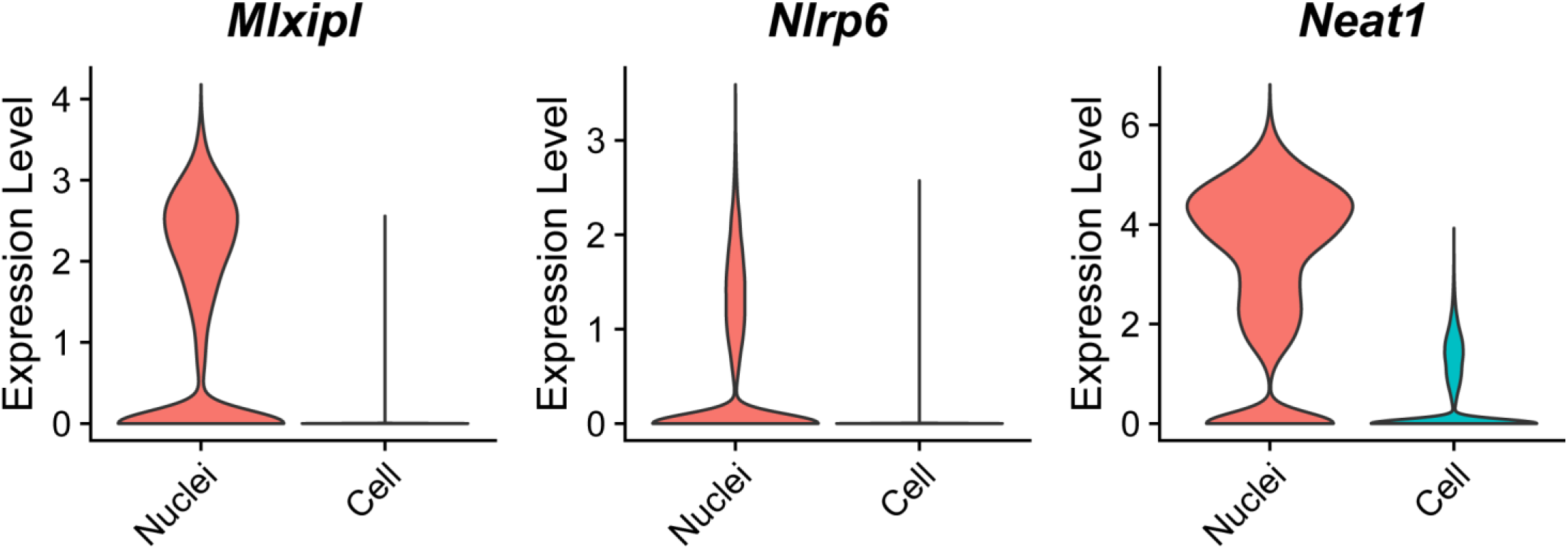
Single nuclei and single cell expression distribution of nuclear biased genes *Mxlipl*, *Nlrp6*, and *Neat1*. Nuclei expression was determined in our dataset, whole cell expression was determined by Xiong et al. (2019) (GSE129516).

**Figure S2.**
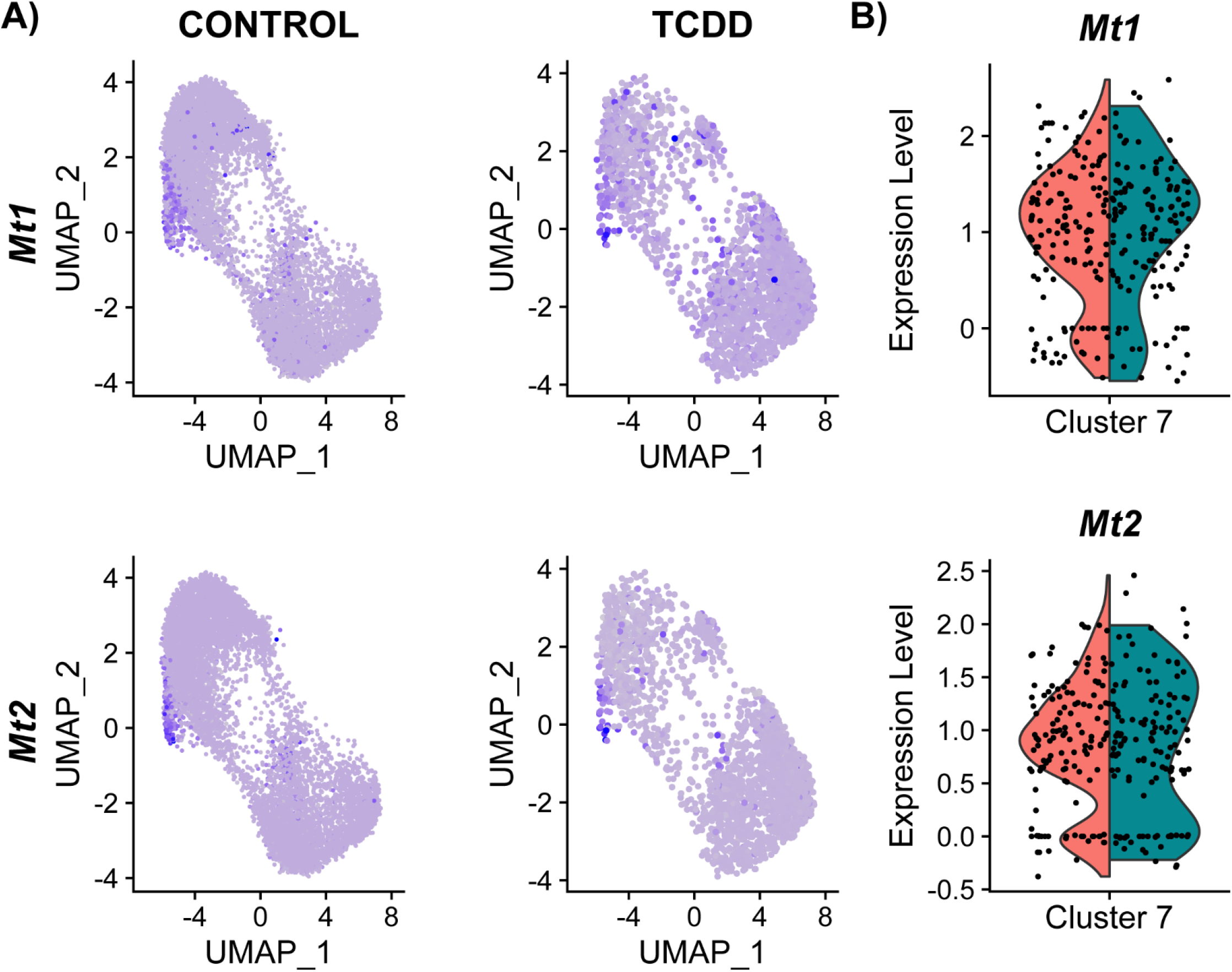
Metallothionein (*Mt1* and *Mt2*) enrichment hepatocytes is visualized using (A) UMAP and (B) violin plots for cluster “7” in control (rose) and TCDD (dark teal). The results demonstrate an absence of differential expression between the groups.

**Figure S3.**
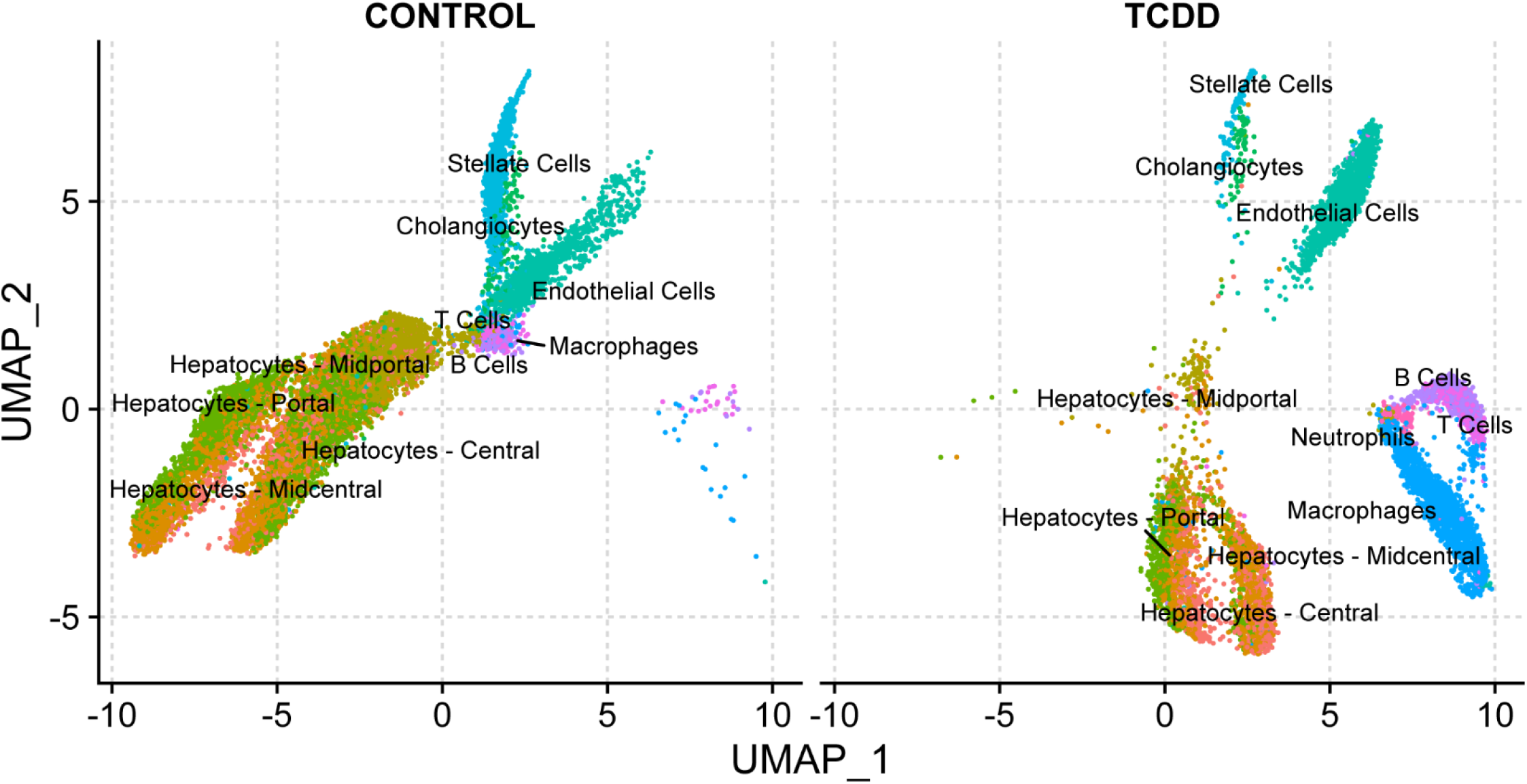
Uniform Manifold Approximation and Projection (UMAP) visualization of nuclei isolated from vehicle and TCDD-treated liver samples clustered based on normalized enrichment score similarity for 6,513 gene sets.

## References

Alvarez, M., Rahmani, E., Jew, B., Garske, K. M., Miao, Z., Benhammou, J. N., Ye, C. J., Pisegna, J. R., Pietiläinen, K. H., Halperin, E. and Pajukanta, P. (2019). Enhancing droplet-based single-nucleus RNA-seq resolution using the semi-supervised machine learning classifier DIEM. bioRxiv, 786285.

Anundi, I., Lahteenmaki, T., Rundgren, M., Moldeus, P. and Lindros, K. O. (1993). Zonation of acetaminophen metabolism and cytochrome P450 2E1-mediated toxicity studied in isolated periportal and perivenous hepatocytes. Biochemical pharmacology 45(6), 1251–9.

Bahar Halpern, K., Caspi, I., Lemze, D., Levy, M., Landen, S., Elinav, E., Ulitsky, I. and Itzkovitz, S. (2015). Nuclear Retention of mRNA in Mammalian Tissues. Cell reports 13(12), 2653–62.

Bakken, T. E., Hodge, R. D., Miller, J. A., Yao, Z., Nguyen, T. N., Aevermann, B., Barkan, E., Bertagnolli, D., Casper, T., Dee, N., Garren, E., Goldy, J., Graybuck, L. T., Kroll, M., Lasken, R. S., Lathia, K., Parry, S., Rimorin, C., Scheuermann, R. H., Schork, N. J., Shehata, S. I., Tieu, M., Phillips, J. W., Bernard, A., Smith, K. A., Zeng, H., Lein, E. S. and Tasic, B. (2018). Single-nucleus and single-cell transcriptomes compared in matched cortical cell types. PloS one 13(12), e0209648.

Ben-Moshe, S., Shapira, Y., Moor, A. E., Manco, R., Veg, T., Bahar Halpern, K. and Itzkovitz, S. (2019). Spatial sorting enables comprehensive characterization of liver zonation. Nature metabolism 1(9), 899–911.

Browaeys, R., Saelens, W. and Saeys, Y. (2020). NicheNet: modeling intercellular communication by linking ligands to target genes. Nature methods 17(2), 159–162.

Butler, A., Hoffman, P., Smibert, P., Papalexi, E. and Satija, R. (2018). Integrating single-cell transcriptomic data across different conditions, technologies, and species. Nature biotechnology 36(5), 411–420.

Cave, M., Appana, S., Patel, M., Falkner, K. C., McClain, C. J. and Brock, G. (2010). Polychlorinated biphenyls, lead, and mercury are associated with liver disease in American adults: NHANES 2003-2004. Environmental health perspectives 118(12), 1735–42.

Digernes, V. and Bolund, L. (1979). The ploidy classes of adult mouse liver cells. A methodological study with flow cytometry and cell sorting. Virchows Archiv. B, Cell pathology including molecular pathology 32(1), 1–10.

Dong, X., Liu, J., Xu, Y. and Cao, H. (2019). Role of macrophages in experimental liver injury and repair in mice. Experimental and therapeutic medicine 17(5), 3835–3847.

Fader, K. A., Nault, R., Doskey, C. M., Fling, R. R. and Zacharewski, T. R. (2019). 2,3,7,8-Tetrachlorodibenzo-p-dioxin abolishes circadian regulation of hepatic metabolic activity in mice. Scientific reports 9(1), 6514.

Fader, K. A., Nault, R., Kirby, M. P., Markous, G., Matthews, J. and Zacharewski, T. R. (2017a). Convergence of hepcidin deficiency, systemic iron overloading, heme accumulation, and REV-ERBalpha/beta activation in aryl hydrocarbon receptor-elicited hepatotoxicity. Toxicol Appl Pharmacol 321, 1–17.

Fader, K. A., Nault, R., Zhang, C., Kumagai, K., Harkema, J. R. and Zacharewski, T. R. (2017b). 2,3,7,8-Tetrachlorodibenzo-p-dioxin (TCDD)-elicited effects on bile acid homeostasis: Alterations in biosynthesis, enterohepatic circulation, and microbial metabolism. Scientific reports 7(1), 5921.

Franzén, O., Gan, L.-M. and Björkegren, J. L. M. (2019). PanglaoDB: a web server for exploration of mouse and human single-cell RNA sequencing data. Database 2019.

Gu, X. and Manautou, J. E. (2012). Molecular mechanisms underlying chemical liver injury. Expert reviews in molecular medicine 14, e4.

Habib, N., Avraham-Davidi, I., Basu, A., Burks, T., Shekhar, K., Hofree, M., Choudhury, S. R., Aguet, F., Gelfand, E., Ardlie, K., Weitz, D. A., Rozenblatt-Rosen, O., Zhang, F. and Regev, A. (2017). Massively parallel single-nucleus RNA-seq with DroNc-seq. Nature methods 14(10), 955–958.

Habib, N., Li, Y., Heidenreich, M., Swiech, L., Avraham-Davidi, I., Trombetta, J. J., Hession, C., Zhang, F. and Regev, A. (2016). Div-Seq: Single-nucleus RNA-Seq reveals dynamics of rare adult newborn neurons. Science 353(6302), 925–8.

Halpern, K. B., Shenhav, R., Massalha, H., Toth, B., Egozi, A., Massasa, E. E., Medgalia, C., David, E., Giladi, A., Moor, A. E., Porat, Z., Amit, I. and Itzkovitz, S. (2018). Paired-cell sequencing enables spatial gene expression mapping of liver endothelial cells. Nature biotechnology.

Halpern, K. B., Shenhav, R., Matcovitch-Natan, O., Toth, B., Lemze, D., Golan, M., Massasa, E. E., Baydatch, S., Landen, S., Moor, A. E., Brandis, A., Giladi, A., Avihail, A. S., David, E., Amit, I. and Itzkovitz, S. (2017). Single-cell spatial reconstruction reveals global division of labour in the mammalian liver. Nature 542(7641), 352–356.

Hanzelmann, S., Castelo, R. and Guinney, J. (2013). GSVA: gene set variation analysis for microarray and RNA-seq data. BMC bioinformatics 14, 7.

Heindel, J. J., Blumberg, B., Cave, M., Machtinger, R., Mantovani, A., Mendez, M. A., Nadal, A., Palanza, P., Panzica, G., Sargis, R., Vandenberg, L. N. and Vom Saal, F. (2017). Metabolism disrupting chemicals and metabolic disorders. Reprod Toxicol 68, 3–33.

Henriksen, G. L., Ketchum, N. S., Michalek, J. E. and Swaby, J. A. (1997). Serum Dioxin and Diabetes Mellitus in Veterans of Operation Ranch Hand. Epidemiology 8(3), 252–258.

Hornberg, J. J., Laursen, M., Brenden, N., Persson, M., Thougaard, A. V., Toft, D. B. and Mow, T. (2014). Exploratory toxicology as an integrated part of drug discovery. Part I: Why and how. Drug discovery today 19(8), 1131–1136.

Iacono, G., Massoni-Badosa, R. and Heyn, H. (2019). Single-cell transcriptomics unveils gene regulatory network plasticity. Genome biology 20(1), 110.

Jackson, D. P., Li, H., Mitchell, K. A., Joshi, A. D. and Elferink, C. J. (2014). Ah receptor-mediated suppression of liver regeneration through NC-XRE-driven p21Cip1 expression. Molecular pharmacology 85(4), 533–41.

Jimenez-Castro, M. B., Cornide-Petronio, M. E., Gracia-Sancho, J., Casillas-Ramirez, A. and Peralta, C. (2019). Mitogen Activated Protein Kinases in Steatotic and Non-Steatotic Livers Submitted to Ischemia-Reperfusion. International journal of molecular sciences 20(7).

Kaiser, J. P., Lipscomb, J. C. and Wesselkamper, S. C. (2012). Putative mechanisms of environmental chemical-induced steatosis. International journal of toxicology 31(6), 551–63.

Katayama, A., Nakatsuka, A., Eguchi, J., Murakami, K., Teshigawara, S., Kanzaki, M., Nunoue, T., Hida, K., Wada, N., Yasunaka, T., Ikeda, F., Takaki, A., Yamamoto, K., Kiyonari, H., Makino, H. and Wada, J. (2015). Beneficial impact of Gpnmb and its significance as a biomarker in nonalcoholic steatohepatitis. Scientific reports 5, 16920.

Krishnaswami, S. R., Grindberg, R. V., Novotny, M., Venepally, P., Lacar, B., Bhutani, K., Linker, S. B., Pham, S., Erwin, J. A., Miller, J. A., Hodge, R., McCarthy, J. K., Kelder, M., McCorrison, J., Aevermann, B. D., Fuertes, F. D., Scheuermann, R. H., Lee, J., Lein, E. S., Schork, N., McConnell, M. J., Gage, F. H. and Lasken, R. S. (2016). Using single nuclei for RNA-seq to capture the transcriptome of postmortem neurons. Nature protocols 11(3), 499–524.

Kucera, M., Isserlin, R., Arkhangorodsky, A. and Bader, G. D. (2016). AutoAnnotate: A Cytoscape app for summarizing networks with semantic annotations. F1000Research 5, 1717.

Lake, B. B., Chen, S., Sos, B. C., Fan, J., Kaeser, G. E., Yung, Y. C., Duong, T. E., Gao, D., Chun, J., Kharchenko, P. V. and Zhang, K. (2018). Integrative single-cell analysis of transcriptional and epigenetic states in the human adult brain. Nature biotechnology 36(1), 70–80.

MacParland, S. A., Liu, J. C., Ma, X. Z., Innes, B. T., Bartczak, A. M., Gage, B. K., Manuel, J., Khuu, N., Echeverri, J., Linares, I., Gupta, R., Cheng, M. L., Liu, L. Y., Camat, D., Chung, S. W., Seliga, R. K., Shao, Z., Lee, E., Ogawa, S., Ogawa, M., Wilson, M. D., Fish, J. E., Selzner, M., Ghanekar, A., Grant, D., Greig, P., Sapisochin, G., Selzner, N., Winegarden, N., Adeyi, O., Keller, G., Bader, G. D. and McGilvray, I. D. (2018). Single cell RNA sequencing of human liver reveals distinct intrahepatic macrophage populations. Nat Commun 9(1), 4383.

Masyuk, T. V., Huang, B. Q., Ward, C. J., Masyuk, A. I., Yuan, D., Splinter, P. L., Punyashthiti, R., Ritman, E. L., Torres, V. E., Harris, P. C. and LaRusso, N. F. (2003). Defects in cholangiocyte fibrocystin expression and ciliary structure in the PCK rat. Gastroenterology 125(5), 1303–10.

Merico, D., Isserlin, R., Stueker, O., Emili, A. and Bader, G. D. (2010). Enrichment map: a network-based method for gene-set enrichment visualization and interpretation. PloS one 5(11), e13984.

Moreno-Marin, N., Merino, J. M., Alvarez-Barrientos, A., Patel, D. P., Takahashi, S., Gonzalez-Sancho, J. M., Gandolfo, P., Rios, R. M., Munoz, A., Gonzalez, F. J. and Fernandez-Salguero, P. M. (2018). Aryl Hydrocarbon Receptor Promotes Liver Polyploidization and Inhibits PI3K, ERK, and Wnt/beta-Catenin Signaling. iScience 4, 44–63.

Nault, R., Fader, K. A., Ammendolia, D. A., Dornbos, P., Potter, D., Sharratt, B., Kumagai, K., Harkema, J. R., Lunt, S. Y., Matthews, J. and Zacharewski, T. (2016). Dose-Dependent Metabolic Reprogramming and Differential Gene Expression in TCDD-Elicited Hepatic Fibrosis. Toxicol Sci 154(2), 253–266.

Nault, R., Fader, K. A., Harkema, J. R. and Zacharewski, T. (2017). Loss of liver-specific and sexually dimorphic gene expression by aryl hydrocarbon receptor activation in C57BL/6 mice. PloS one 12(9), e0184842.

Nault, R., Fader, K. A. and Zacharewski, T. (2015). RNA-Seq versus oligonucleotide array assessment of dose-dependent TCDD-elicited hepatic gene expression in mice. BMC genomics 16(1), 373.

Nonaka, H., Tanaka, M., Suzuki, K. and Miyajima, A. (2007). Development of murine hepatic sinusoidal endothelial cells characterized by the expression of hyaluronan receptors. Developmental Dynamics 236(8), 2258–2267.

Ohguchi, H., Tanaka, T., Uchida, A., Magoori, K., Kudo, H., Kim, I., Daigo, K., Sakakibara, I., Okamura, M., Harigae, H., Sasaki, T., Osborne, T. F., Gonzalez, F. J., Hamakubo, T., Kodama, T. and Sakai, J. (2008). Hepatocyte nuclear factor 4alpha contributes to thyroid hormone homeostasis by cooperatively regulating the type 1 iodothyronine deiodinase gene with GATA4 and Kruppel-like transcription factor 9. Molecular and cellular biology 28(12), 3917–31.

Pierre, S., Chevallier, A., Teixeira-Clerc, F., Ambolet-Camoit, A., Bui, L. C., Bats, A. S., Fournet, J. C., Fernandez-Salguero, P., Aggerbeck, M., Lotersztajn, S., Barouki, R. and Coumoul, X. (2014). Aryl hydrocarbon receptor-dependent induction of liver fibrosis by dioxin. Toxicol Sci 137(1), 114–24.

Santostefano, M. J., Richardson, V. M., Walker, N. J., Blanton, J., Lindros, K. O., Lucier, G. W., Alcasey, S. K. and Birnbaum, L. S. (1999). Dose-dependent localization of TCDD in isolated centrilobular and periportal hepatocytes. Toxicol Sci 52(1), 9–19.

Smith, X., Taylor, A. and Rudd, C. E. (2016). T-cell immune adaptor SKAP1 regulates the induction of collagen-induced arthritis in mice. Immunology letters 176, 122–7.

Street, K., Risso, D., Fletcher, R. B., Das, D., Ngai, J., Yosef, N., Purdom, E. and Dudoit, S. (2018). Slingshot: cell lineage and pseudotime inference for single-cell transcriptomics. BMC genomics 19(1), 477.

Stuart, T., Butler, A., Hoffman, P., Hafemeister, C., Papalexi, E., Mauck, W. M., 3rd, Hao, Y., Stoeckius, M., Smibert, P. and Satija, R. (2019). Comprehensive Integration of Single-Cell Data. Cell 177(7), 1888–1902 e21.

Su, X., Shi, Y., Zou, X., Lu, Z. N., Xie, G., Yang, J. Y. H., Wu, C. C., Cui, X. F., He, K. Y., Luo, Q., Qu, Y. L., Wang, N., Wang, L. and Han, Z. G. (2017). Single-cell RNA-Seq analysis reveals dynamic trajectories during mouse liver development. BMC genomics 18(1), 946.

Taylor, K. W., Novak, R. F., Anderson, H. A., Birnbaum, L. S., Blystone, C., Devito, M., Jacobs, D., Kohrle, J., Lee, D. H., Rylander, L., Rignell-Hydbom, A., Tornero-Velez, R., Turyk, M. E., Boyles, A. L., Thayer, K. A. and Lind, L. (2013). Evaluation of the association between persistent organic pollutants (POPs) and diabetes in epidemiological studies: a national toxicology program workshop review. Environmental health perspectives 121(7), 774–83.

Xiong, X., Kuang, H., Ansari, S., Liu, T., Gong, J., Wang, S., Zhao, X. Y., Ji, Y., Li, C., Guo, L., Zhou, L., Chen, Z., Leon-Mimila, P., Chung, M. T., Kurabayashi, K., Opp, J., Campos-Perez, F., Villamil-Ramirez, H., Canizales-Quinteros, S., Lyons, R., Lumeng, C. N., Zhou, B., Qi, L., Huertas-Vazquez, A., Lusis, A. J., Xu, X. Z. S., Li, S., Yu, Y., Li, J. Z. and Lin, J. D. (2019). Landscape of Intercellular Crosstalk in Healthy and NASH Liver Revealed by Single-Cell Secretome Gene Analysis. Mol Cell 75(3), 644–660 e5.

Yamaguchi, M. and Hankinson, O. (2018). 2,3,7,8Tetrachlorodibenzopdioxin suppresses the growth of human liver cancer HepG2 cells in vitro: Involvement of cell signaling factors. International journal of oncology 53(4), 1657–1666.

Yang, S., Corbett, S. E., Koga, Y., Wang, Z., Johnson, W. E., Yajima, M. and Campbell, J. D. (2019). Decontamination of ambient RNA in single-cell RNA-seq with DecontX. bioRxiv, 704015.

Yang, S. and Liu, G. (2017). Targeting the Ras/Raf/MEK/ERK pathway in hepatocellular carcinoma. Oncology letters 13(3), 1041–1047.

Yorita Christensen, K. L., Carrico, C. K., Sanyal, A. J. and Gennings, C. (2013). Multiple classes of environmental chemicals are associated with liver disease: NHANES 2003-2004. International journal of hygiene and environmental health 216(6), 703–9.

Young, M. D. and Behjati, S. (2020). SoupX removes ambient RNA contamination from droplet based single-cell RNA sequencing data. bioRxiv, 303727.

Zeng, W., Jiang, S., Kong, X., El-Ali, N., Ball, A. R., Jr., Ma, C. I., Hashimoto, N., Yokomori, K. and Mortazavi, A. (2016). Single-nucleus RNA-seq of differentiating human myoblasts reveals the extent of fate heterogeneity. Nucleic acids research 44(21), e158.

Zhang, Z., Cotta, C. V., Stephan, R. P., deGuzman, C. G. and Klug, C. A. (2003). Enforced expression of EBF in hematopoietic stem cells restricts lymphopoiesis to the B cell lineage. The EMBO journal 22(18), 4759–69.

